# Host-Microbe Interactions in Manifestation of Tuberculosis: A System Biology Study in Implicated Compartments

**DOI:** 10.1101/2020.12.06.413617

**Authors:** Sharebiani Hiva, Abbasnia Shadi, Soleimanpour Saman, Rezaee Sar

## Abstract

*Mycobacterium tuberculosis* (*Mtb*) has been a dilemma for over a century. Thus the bacteria-host interactions seem to be implicated in the manifestation of the disease. Here, the behavioral activities of the *Mtb* and host responses were evaluated in this system biology analyses, according to the compartmental immune responses in the lung and local lymph node. Differential expression analyses were conducted between tuberculosis (TB) and the healthy group in the aforementioned compartments, to identify the hub genes and functional gene ontology (GO) terms, using KEGG, Enrichr and DAVID databases.

The different phases of immune responses against *Mtb* occur in three compartments, lung, local lymph nodes and blood. Due to the occurrence of hypoxia within granuloma in the lung, angiogenesis was increased despite the HIF1-α down-regulation via inhibition of EP300 and HDAC1. Proliferation by MYC, CDK2 and NF-κB pathways activated in the granuloma, while at the same time apoptosis was induced by P53 activation, and PI3K/Akt inhibited P53 in the lymph node. Furthermore, DNA damages suppressed by the over-expression of BRCA1, CDK1 and BCR/ABL in the lymph node, as well as FBXO6, CDK2 and CDC5A in both compartments. In the lymph node, RTK (EGFR) and calmodulin, the consequent NFAT formation and Erk/MAPK pathway down-regulated and suppressed Th1 cell activation and differentiation. Inflammation was induced in both compartments, but the antigen (Ag) presentation was suppressed through the XPO1 suppression and ubiquitination. More studies in *Mtb*-host interactions are needed to specify the effective mechanisms for reducing this re-emerging life-threatening disease.

**Importance:** Tuberculosis (TB) is one of the most widespread reemerging infectious diseases in the world, which has remained a global health problem. Approximately, 10 million people are infected with *Mycobacterium tuberculosis* (*Mtb*), causing 1.2 million deaths every year. Therefore, interactions between the host and the pathogen in *Mtb* infection are a major challenge for the control of the disease. Typically, there are thousands of genes and ten times more interactions between any stages of the conflicts. This urged us to bring “systemic approaches” for a better understanding of such highly orchestrated systems. A holistic view of the *Mtb*-host interaction paves the way for a higher insight into the biology of the organism, as well as rationale solutions for the design of therapeutic agents. This study specifies the nominated disease-related genes and related signaling pathways in the pathogenesis of TB in two different compartments, lung and lymph node.

## Introduction

Tuberculosis (TB) is one of the most widespread reemerging infectious diseases in the world, which has remained a global health problem. According to the World Health Organization (WHO) reports, approximately 10 million (ranging from 9 to 11.1 million) people are infected with *Mycobacterium tuberculosis* (*Mtb*), causing 1.2 million deaths (ranging from 1.1 to 1.3 million) every year [1]. *Mtb*, as the causative agent of TB and interaction factor between the host and the pathogen in *Mtb* infection, is a major challenge for the control of the disease [2].

Upon inhalation of *Mtb*, the bacteria are taken up by alveolar macrophages and DCs, and are finally led to the formation of the granuloma [3]. The granuloma is a complex compartment of *Mtb* and host mononuclear cells with a fibrin ring, leading the host response to *Mtb* colonization in order to protect the surrounding tissue [3]. To produce a protective response, the host infected macrophages or DCs, engulfing the *Mtb*, migrate into the T cell zone in the regional lymph nodes, where specific *Mtb* CD4^+^ T cells recognize *Mtb* antigens in a Th1-inducing micro-environment [4]. The process results in the activation of *Mtb*-specific T lymphocytes, which are then migrating to the sites of *Mtb* replication [4]. On the other hand, the development and progression of TB result in the formation of a caseous granuloma that cavitates and spreads *Mtb* [5]. Of note, this form of granuloma is an immune-pathologic manifestation of inappropriate host responses in favor of *Mtb*. Although important advances have been made in the control of some acute infections, using interventions that target a single biological focal point is more difficult to anticipate the impact of a targeted intervention on the complex biology of persistent infection [5]. Nowadays, it is well known that the development and progression of infectious diseases depend on the molecular interactions between the microbe and the host [6]. However, the identification of such molecules in different phases of infection, from elimination to the manifestation of a disease, is a great challenge [6]. Typically, there are thousands of genes and ten times more interactions between any stages of the conflicts. This urged us to bring “systemic approaches” for a better understanding of such highly orchestrated systems, as well as manipulating the system accordingly, if necessary [7]. The methodological point of view, introducing a reliable scientific method for interpretation of the complex interactions between two intelligent organisms, fighting for the survival of their species is a dilemma, particularly to find the precise target for the intervention, in favor of the host [6]. A holistic view of the *Mtb* and its interaction with the host, provide a basis for higher insight into the biology of the organism, as well as rationale solutions for the design of therapeutic agents. Mathematical medicine has recently focused to resolve this problem using algorithms, data mining, game theory, and complex system biology, as well as the robotic calculations [8]. In the current study, to pave a path for understanding such communications, the *Mtb*-host interactions in two different compartments, such as lung and lymph node, were assessed by the network-based analyses. The results specify the nominated disease-related genes and related signaling pathways in the pathogenesis.

## Materials and Methods

### Gene expression microarray datasets

The gene expression profiles of the caseous human pulmonary TB granulomas and lymph node of the TB patients were obtained from the NCBI Gene Expression Omnibus (GEO) database (Accession no: GSE20050 and GSE63548, respectively). Therefore, as the authors did not any dealing with the human or animal materials, and used the data from deposited free data form GEO database (https://www.ncbi.nlm.nih.gov/geo/) the Ethical Approval for this system biology study is not applicable. Of note, the reference numbers of the experiments with all necessary documents regarding these data were included in these sections

In other words this is a comprehensive system biology analysis on GEO data in different compartments of host immune system to the M. tuberculosis in a comprehensive manner.

I was wondering if you could let me know if I should do something in particular for such

In the first study, the authors used laser capture microdissection (LCM) to dissect out the caseous granulomas from the lung tissues of TB patients, excluding the uninvolved areas. Total RNA was isolated from LCM-derived materials and used for the microarray. As a control, parenchyma from normal lung tissues was prepared in a similar manner. Gene expression was performed, using Affymetrix Human X3P Array (Platform accession no: 1352). In this microarray study, 7 samples were enrolled within two groups, i.e. caseous human pulmonary TB granulomas (*n* = 5) and normal lung (*n* = 2). In the second study, total RNA was extracted from the lymph node tissue samples of the TB patients (*n*=22); for the control group, the RNA samples of lymph node of healthy subjects (*n*=4) and their global transcriptome profiling were considered, using Illumina Human HT-12 V4 expression bead chip (Platform accession no: GPL10558).

### Analysis of Differentially Expressed Genes (DEGs) and Log Fold Change (LogFC)

In both studies, the bioinformatics analysis of differences in gene expression between the two groups, and calculation of the fold change (FC) was assessed using GEO2R, which was employed to normalize the datasets with log2 transformation [9]. The results with an adjusted *p*-value of less than 0.05 were considered statistically significant. Also, the logFC was considered as a criterion to choose the up-regulated (positive logFC) and down-regulated (negative logFC) genes. Finally, the heat map plots, correlation matrix and gene expression profile were generated, using the package of the heat map in R 3.2.5.

### Centrality analysis and Protein-Protein Interaction Networks (PPINs)

Cytoscape software (V3.5.1) was used to create the networks and their analysis. Bisogenet application is a plugin of this software, which was applied for searching the molecular interactions, and displaying the results as a PPIN. All the accessible interaction sources, comprising physical interactions and functional associations were derived from the NCBI, UniProt, DAVID and KEGG databases.

The PPINs were analyzed by this network analyzer application, and the centrality parameters, including the degree, closeness, and betweenness were calculated. The degree of a node is defined as the number of edges, which are interconnected. The top 50 genes with higher scores in three aforesaid parameters were identified and their common genes were selected as the hub genes. In the next step, PPIN of hub genes was generated based on the associated statistical parameters, to be stated later as the centrality indices.

### Hub genes enrichment analysis

The DAVID bioinformatics resource 6.8, was utilized as a web tool to enrich the hub genes in terms of gene ontology (GO) [10]. Furthermore, the significantly enriched KEGG pathway terms were considered, based on the top ten combined scores [11].

### The Tuberculosis Implicated Signaling Network (TISN)

The signaling network of the TB was created, according to the literature reports and the KEGG pathways. The genes implicated in various pathways, including different roles in the pathogenicity of TB were identified and then merged to construct the TISN. The down-regulated genes were ascertained by blue and green colors, and the up-regulated genes were designated as red and yellow colors. Blue and red colors are similar between both tissue granuloma and lymph node, while the green and yellow colors illustrate the changes in the expression of associated genes in the manifestation of TB.

## RESULTS

### Heat map plots; the sample-sample correlation matrix and hub gene expression

The correlation matrix of samples is shown in Figs.1 and 2, in the format of the heat map plot with a range of colors for the granuloma and lymph node, respectively (blue to red showing no difference to the higher correlation, respectively). The correlation matrix was calculated by comparing the expression data, based on the Pearson correlation.

**Figure 1.**
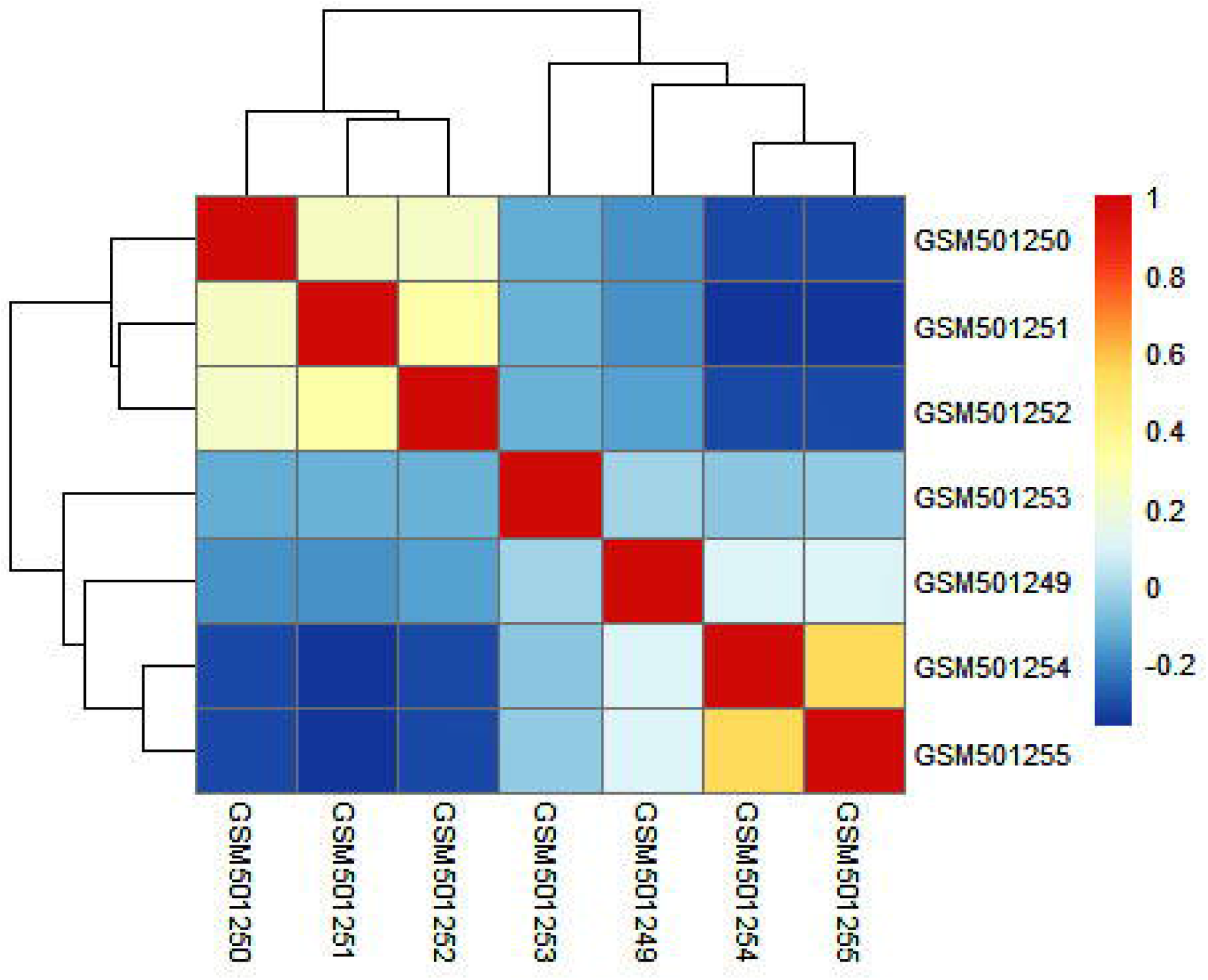
The heat map plot, representing the pairwise correlation across different GSMs of normal and granuloma lung samples. The colors show the relative correlation between 7 samples, as specified in the color key.

**Figure 2.**
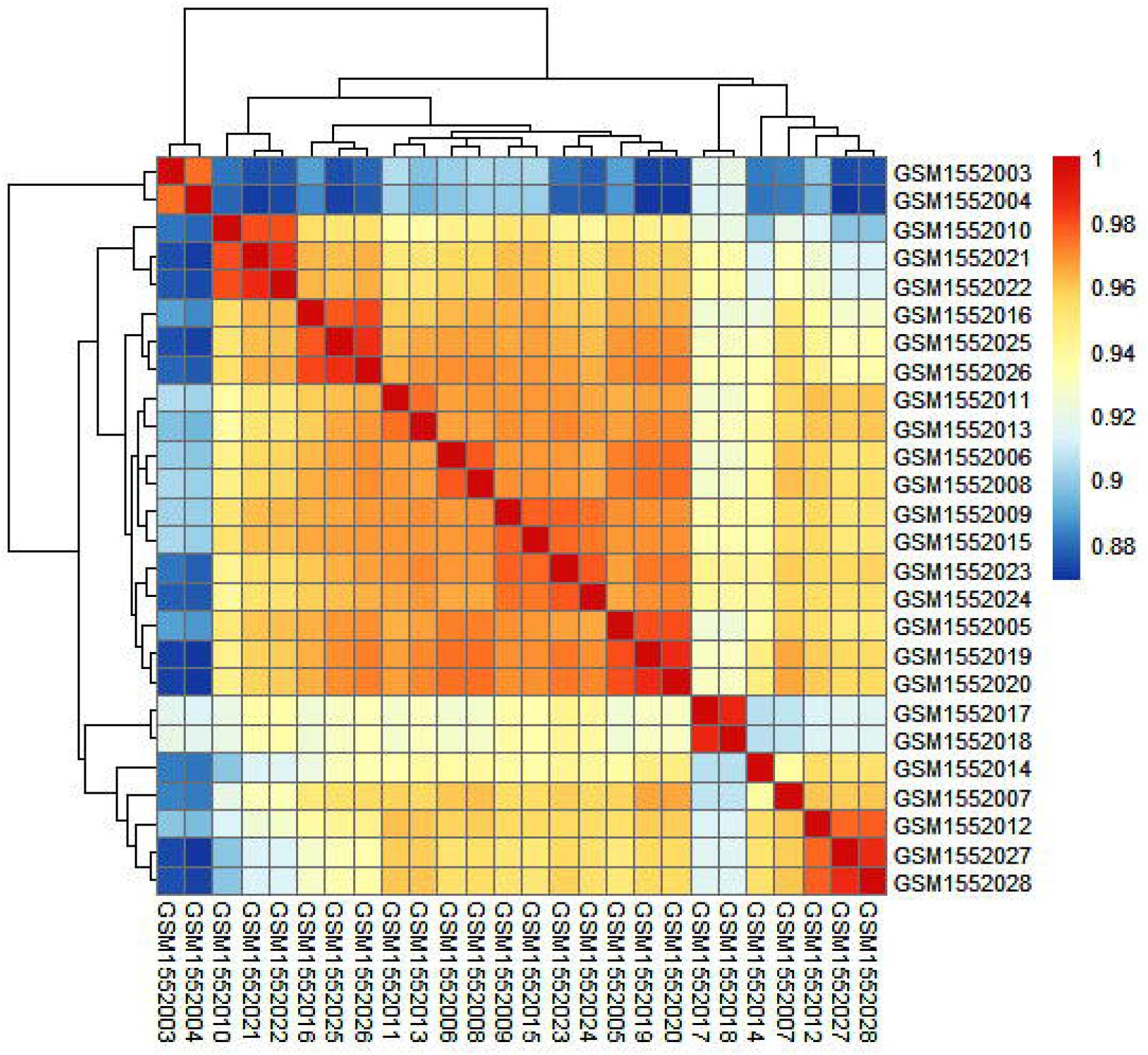
The pairwise sample correlation across different GSMs of normal and TB lymph node samples, represented by the heat map plot. The colors show the relative correlation between 26 samples, as specified in the color key.

The overlap between the gene expression profiles of samples is displayed by a high correlation between the samples. Furthermore, the heat map plot of the hub genes between the two groups of caseous human pulmonary TB granulomas and normal lung was presented in Fig. 3, and the plot of hub genes between TB and normal lymph node depicted in Fig. 4. Thus, the mean of the gene expression was calculated for the samples of each group. The down-regulated genes are shown in green and the up-regulated ones are presented in red.

**Figure 3.**
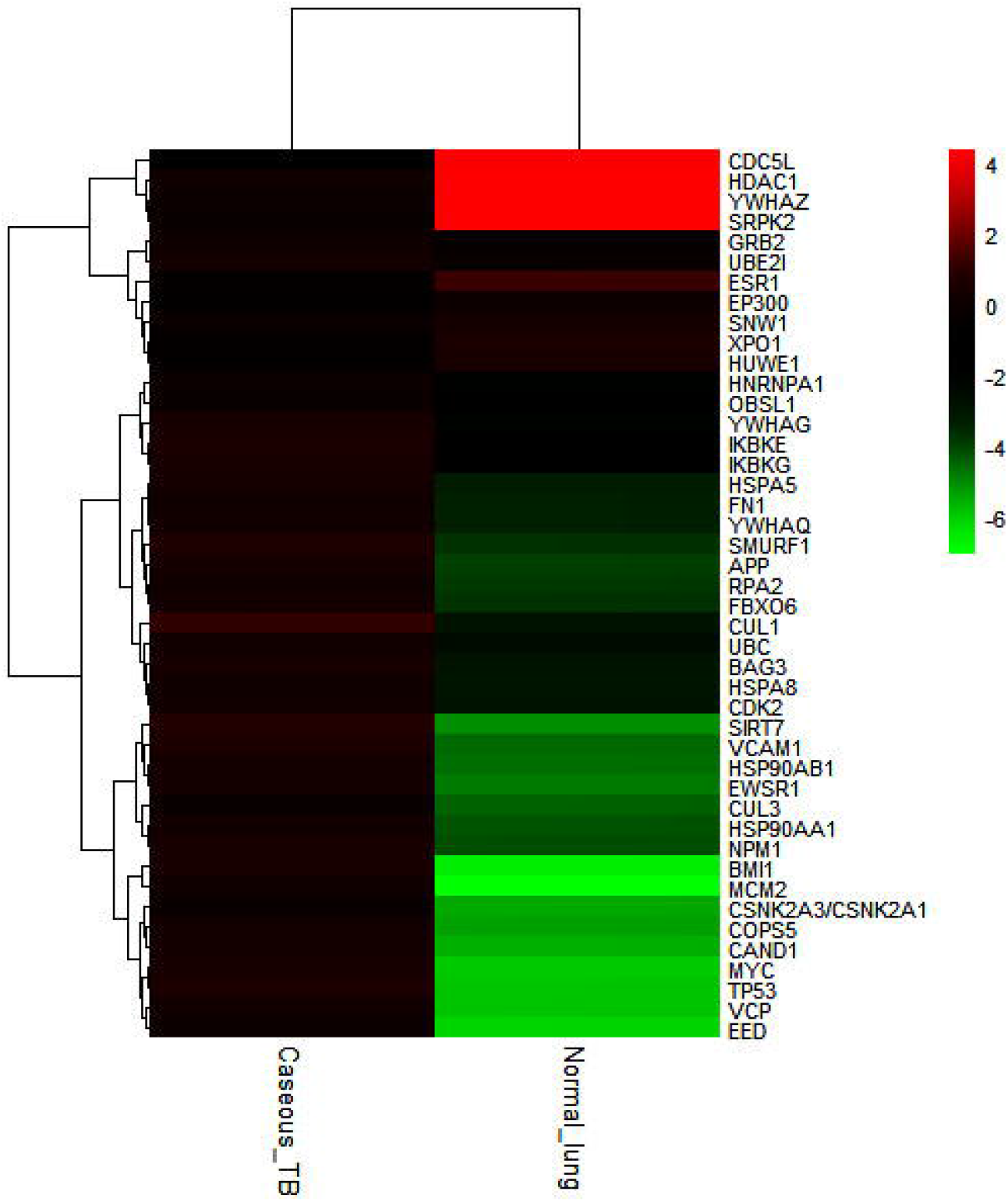
The heat map of the hub genes between two groups of caseous human pulmonary TB granulomas and normal lung. The colors demonstrate the expression level of each gene, as specified in the color key.

**Figure 4.**
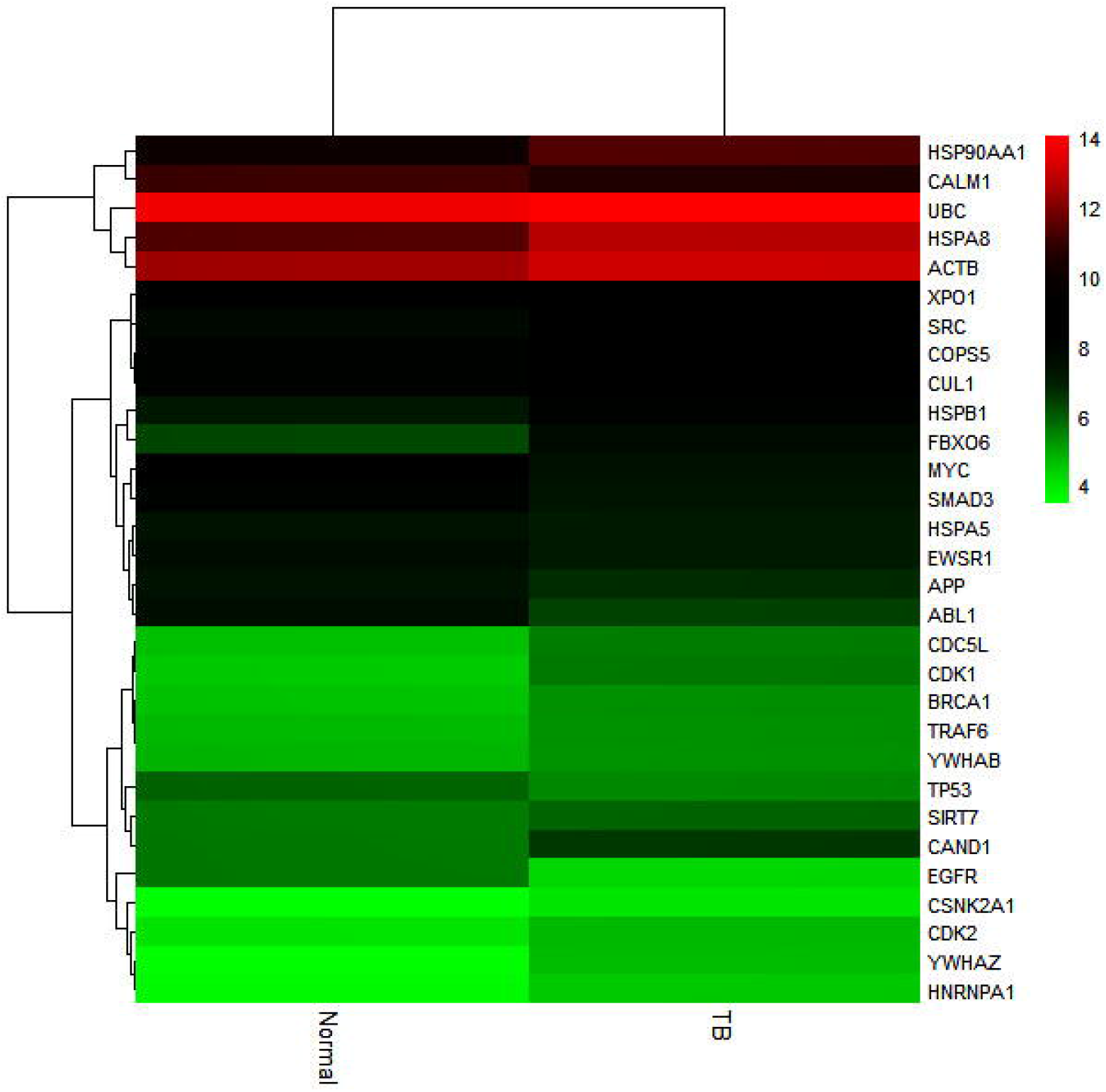
The heat map of the hub genes between two groups of lymph node tissues from healthy individuals (normal) and *Mtb* infected lymph nodes of patients (TB). The colors demonstrate the expression level of each gene, as specified in the color key.

### DEGs and hub gene identification

Based on the criteria of the adjusted *p*-value of less than 0.05, 8151 and 6692 genes with remarkable differential expressions were identified in the granuloma and lymph node studies, respectively. Various measures of the centrality were considered within the graph theory and network analysis to identify the hub genes. As a whole, 45 genes were identified as the hub genes in granuloma study and 30 genes determined as the lymph node hub genes. The up-regulated and down-regulated hub genes are illustrated in Table 1, with the value of the logFC.

### Protein-Protein Interaction Network (PPIN)

The PPINs of hub genes and clusters were generated to display the connectivity and centrality between the identified significant genes, in order to explain the key points in the pathogenesis of TB (Figs. 5 and 6). The variation of hub gene networks consists of 45 differentiated nodes in the caseous human pulmonary TB granulomas in Fig. 5, and also 30 nodes, including the hub genes in the lymph node of TB patients are demonstrated in Fig. 6. The sizes and colors of nodes and edges in all the PPINs were defined by the centrality indices, including the degree, betweenness and closeness.

**Figure 5.**
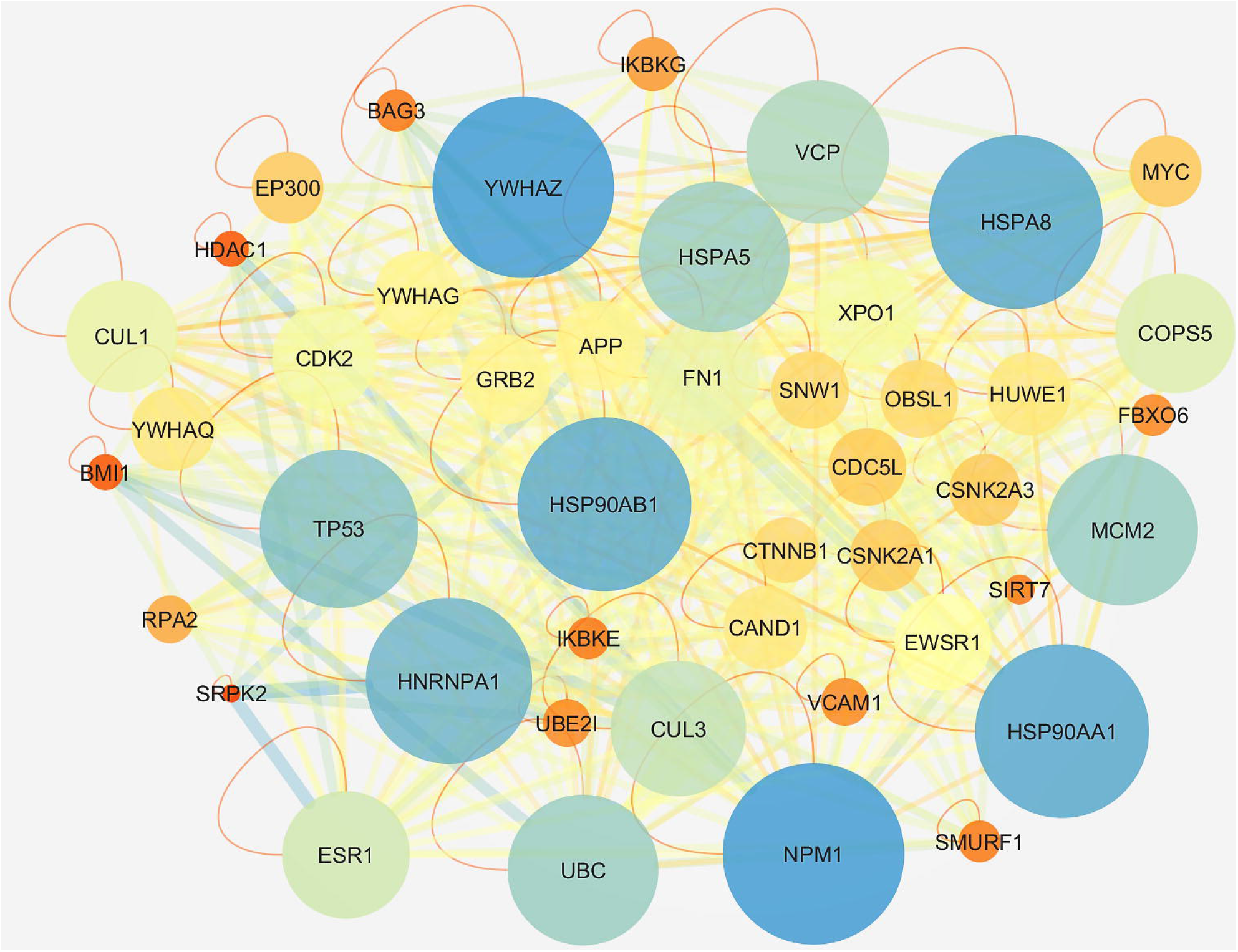
The PPINs of hub genes between two groups of caseous human pulmonary TB granulomas and normal lung. The node size displays the degree of nodes and the node color represents the closeness centrality of nodes, ranging from red (low value) to blue (high value). Also, the thickness of edges and edge color represent betweenness centrality. High values indicated by a thick line and dark color (blue).

**Figure 6.**
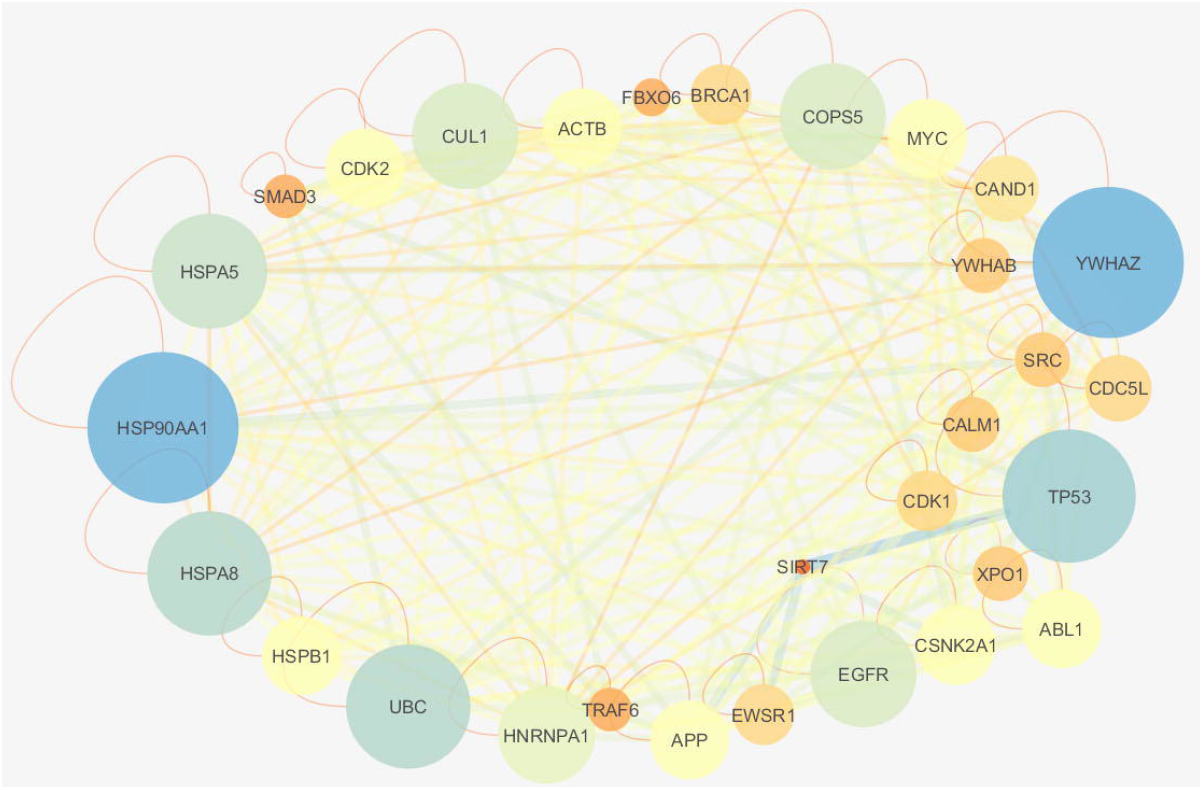
The PPINs of hub genes between two groups of TB and normal lymph nodes. The node size displays the degree of nodes and the node color represents the closeness centrality of nodes, ranging from red (low value) to blue (high value). Also, the thickness of edges and edge color represent betweenness centrality. High values indicated by a thick line and dark color (blue).

### The shared hubs between granuloma and lymph node in TB patients

The Venn diagram in Fig. 7 represents 19 common hub genes between granuloma and lymph node and other hub genes, specific for each compartment (section) such as in the granuloma, monocyte-macrophage, and in the lymph node, dendritic cell-lymphocyte. These commonly expressed genes are illustrated by red and blue colors, which represent the up-regulation and down-regulation, respectively.

**Figure 7.**
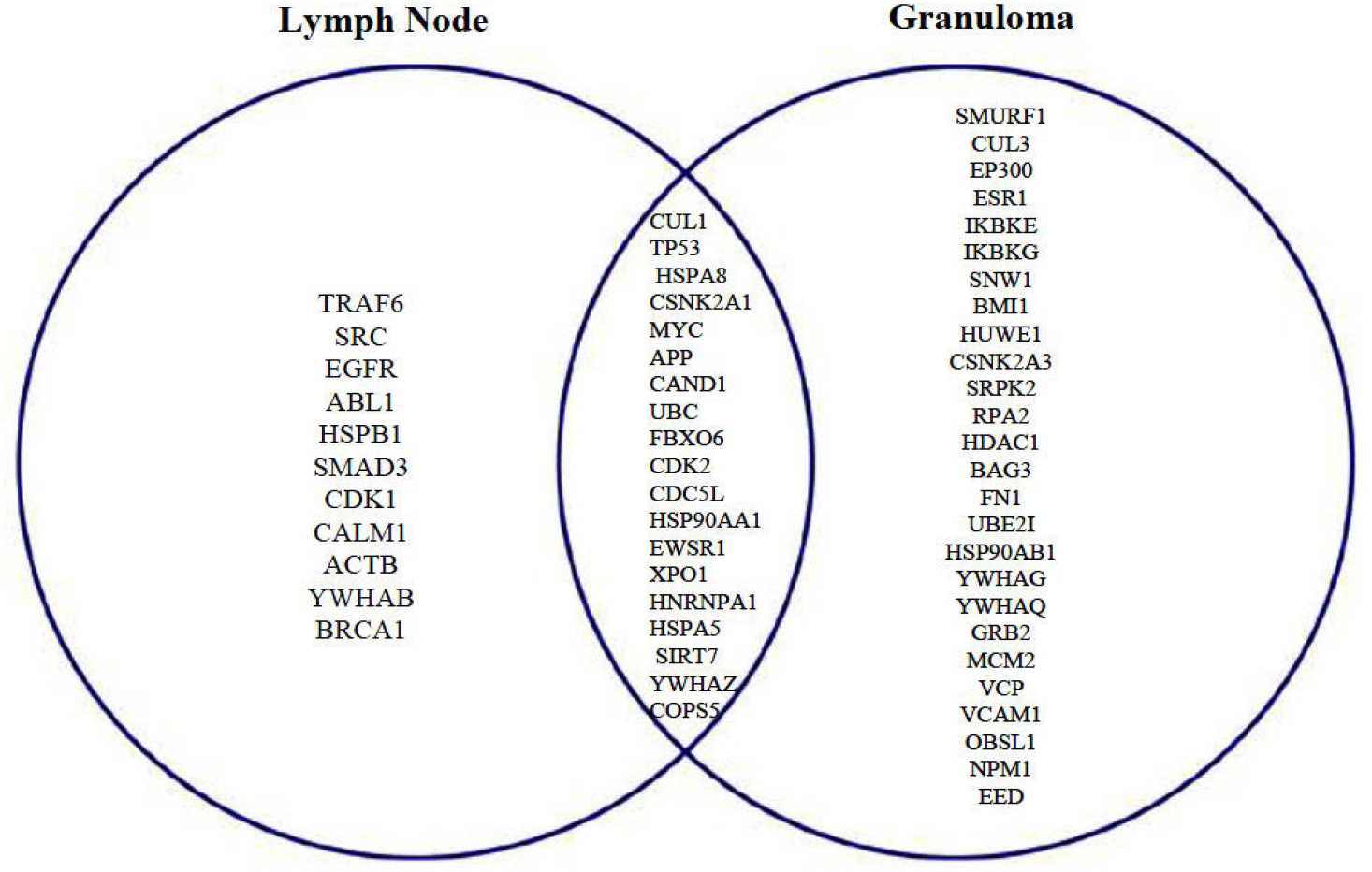
The Venn diagram of lymph node and granuloma hub genes.

### The Tuberculosis Implicated Signaling Network (TISN)

The signaling network and GO of granuloma hub genes are illustrated in Fig. 8 and Table 2, respectively. The pathways in the granuloma signaling network (GSN) could be up-regulated in the diseases, including actin polymerization, anaerobic metabolism, glutaminolysis, fatty acid oxidation and lipid uptake, angiogenesis, cell proliferation, apoptosis and autophagy. While some biological processes, such as antigen (Ag) presentation and pentose phosphate pathway are down-regulated, some processes including inflammation and glucose metabolism are under control by different pathways of inhibition and activation. Two main hubs in this network that might be more implicated in the TB manifestation are MYC and P53, which are common in NF-κB, PI3K-Akt, MAPK, Wnt and focal adhesion pathways. These two up-regulated genes lead to the activation of some processes, such as anaerobic metabolism, glutaminolysis, fatty acid oxidation and lipid uptake, angiogenesis, proliferation, apoptosis and autophagy, as well as inhibition of inflammation. On the other hand, some other up-regulated genes such as UBE2I and IKKs are involved in the activation of the inflammation process. It is noteworthy to state that fibronectin-1 (FN1) and VCAM-1 play key roles in the activation of focal adhesion and PI3K-Akt pathways in this network (Fig. 8).

**Figure 8.**
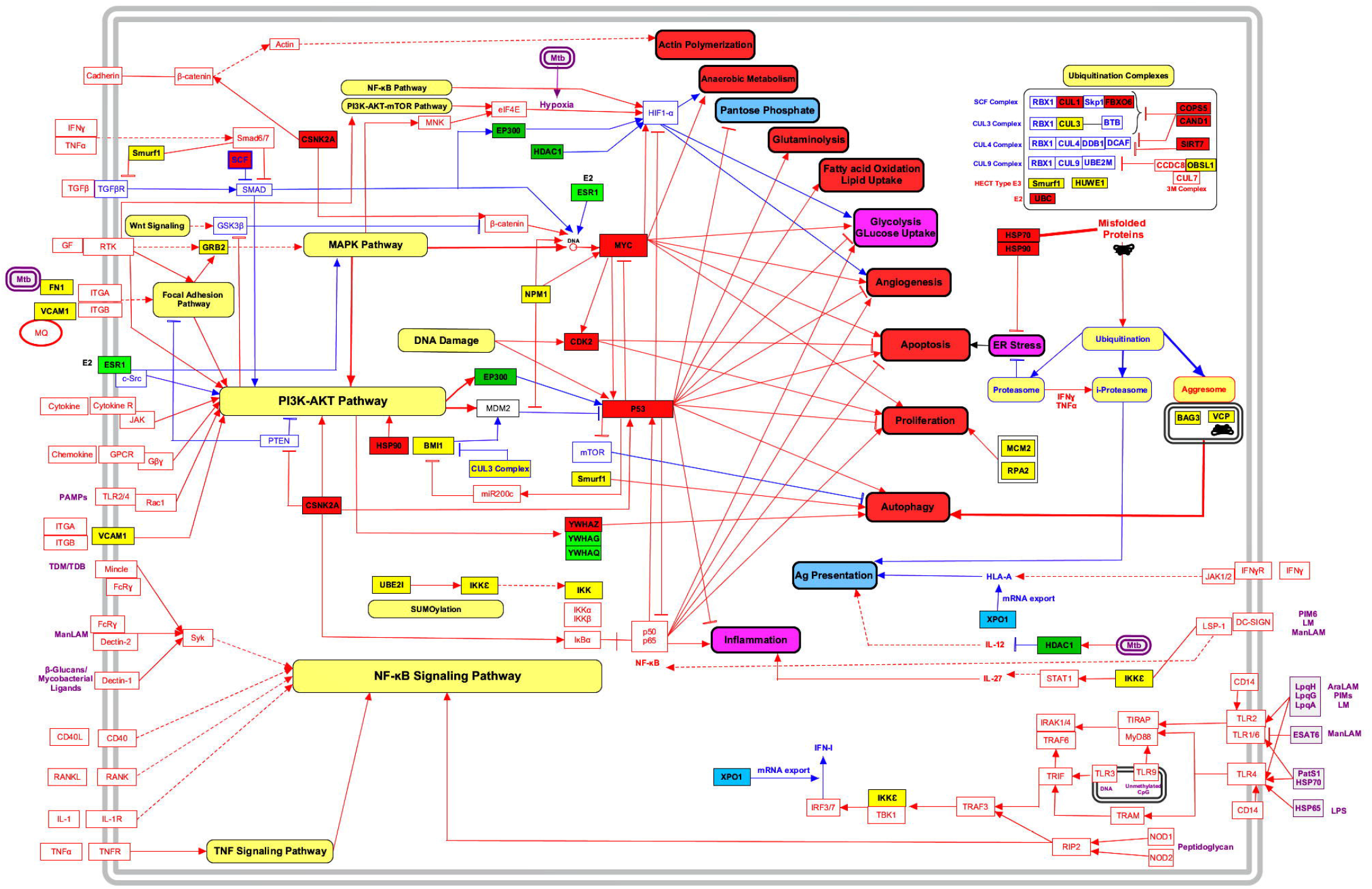
Granuloma Signaling Network (GSN). Blue and green colors show down-regulation, and red and yellow colors are upregulated genes in the granuloma. Green and yellow colors represent genes, which are specific for the granuloma. Red and blue colors represent common genes between granuloma and lymph node. Blue lines are down-regulated pathways and red lines are up-regulated pathways. Solid and dotted arrows demonstrate direct and indirect interactions, respectively.

Activation of SUMOylation in this network is involved in NF-κB pathway activation and induction of inflammation. Moreover, suppression of some proteins in ubiquitin systems such as SCF, CUL3, CUL4 and CUL9 complex, and also their inhibitors, including COPS5, CAND1, OBSL1 and SIRT7 occurs. These processes in the network can induce severe ER stress and high expression of chaperon molecules such as HSPs (Fig. 8).

These three steps in the downstream of ubiquitination, such as the proteasome, immune-proteasome (i-proteasome) and aggresome are important for *Mtb* to be targeted. Therefore, inducing the high expression of BAG3 and VCP in the aggresome complex and expression of P53, YWHAZ and SMURF1 as the activator of autophagy pathways were up-regulated, as an attempt to put pressure on *Mtb*. However, at the same time by such variations in the ubiquitin complex, the processing of the antigen pathway is also inhibited, and consequently, i-proteasome, which is an orchestrated Ag presentation, is suppressed in favor of *Mtb* dissemination. In addition, the down-regulation of XPO1 and HDAC1gene expression decreases the Ag presentation capacity of the infected APCs (Fig. 8).

As a result, there is a low Ag presentation, high cell turnover (severe proliferation and apoptosis at the same time), high-activated cell metabolism in granuloma location or monocyte and macrophage cells in caseous stage, and following to this level of metabolism, oxygen uptake is elevated by angiogenesis in order to provide this level of metabolism in the high turnover cells. Although apoptosis is activated in this network to eliminate the infection, some other survival processes are activated to compensate the host attempt to induce the cell death mechanisms (Fig. 8). Furthermore, the GO and signaling network of lymph node hub genes are represented in Fig. 9 and Table 3. Some pathways, including NF-κB, PI3K-Akt, MAPK, Wnt and focal adhesion pathways, almost such as granuloma, are up-regulated in the lymph node signaling network (LNSN). These pathways in lymph node involved in (i) activation of a group of biological processes such as anaerobic metabolism, pentose phosphate pathway, glucose uptake and glycolysis, inflammation, angiogenesis and proliferation, (ii) inhibition of glutaminolysis, fatty acid oxidation and lipid uptake, autophagy, apoptosis, Ag presentation, phagolysosome formation and TH1/TH17 differentiation, and (iii) control of some processes such as inflammation and actin polymerization by both activator and inhibitor signals. The status of ubiquitination is similar to GSN through inhibition of SCF and CUL4 complex by COPS5, CAND1 and SIRT7. The three next steps other than ubiquitination are down-regulated, and therefore both of the autophagy and Ag presentations are down-regulated. The highest variation in LNSN with GSN decrements in RTK and CALM expression levels, which suppresses phagolysosome formation, Th1/Th17 differentiation and proliferation.

**Figure 9.**
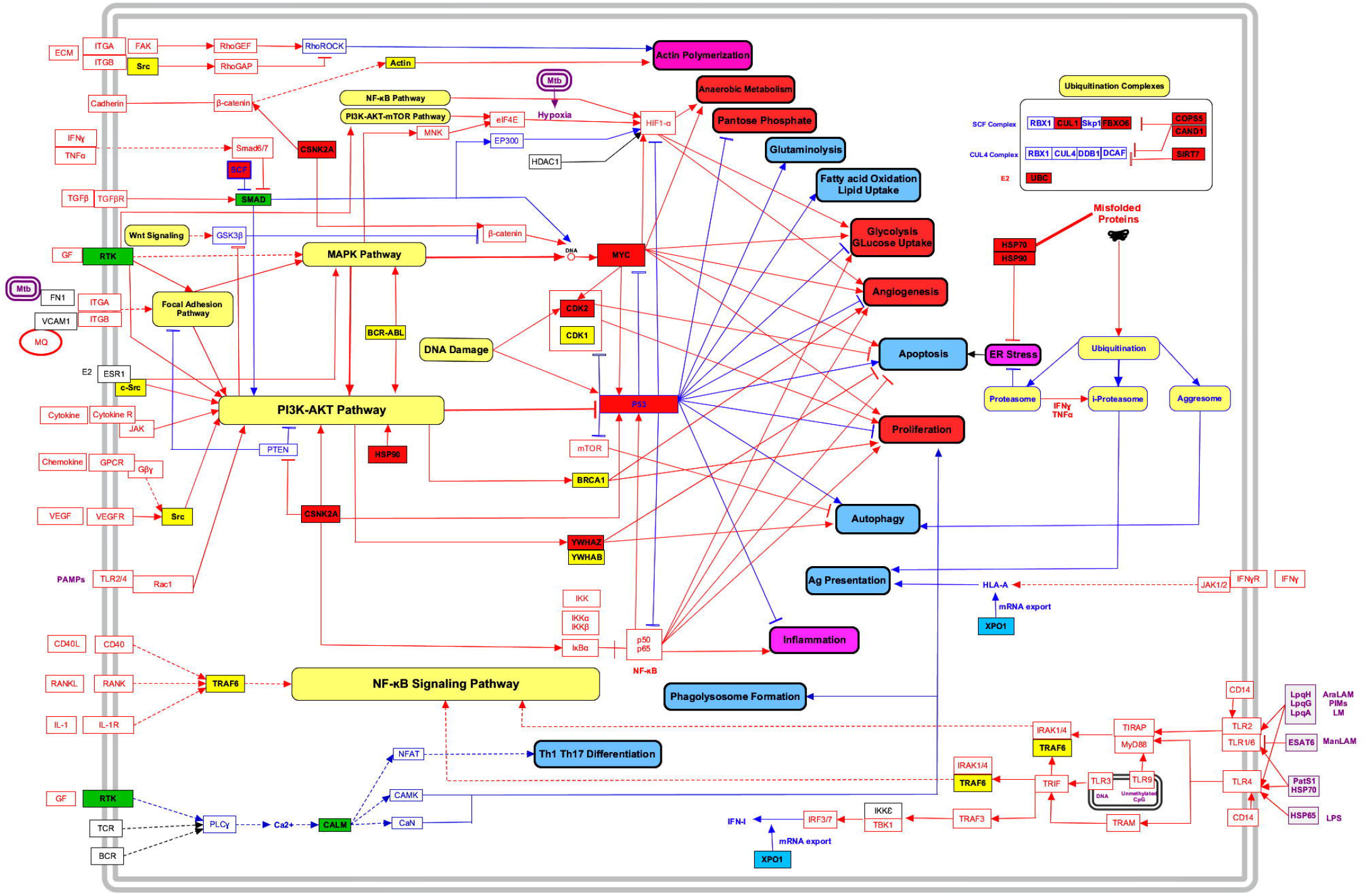
Lymph Node Signaling Network (LNSN). Blue and green colors show down-regulation, and red and yellow colors are up-regulated genes in the granuloma. Green and yellow colors represent genes, specifically for the lymph node. Red and blue colors represent common genes between granuloma and lymph node. Blue lines are down-regulated pathways and red lines are up-regulated pathways. Solid and dotted arrows demonstrate direct and indirect interactions, respectively.

NPM1 is an important molecule in GSN, which acts as an inhibitor of MDM2 that finally activates high expression of P53, following the activation of the PI3K-Akt pathway. However, in LNSN this molecule is not a hub gene and is not expressed in high amounts. Therefore, the P53 suppression in LNSN can be a marker in LNSN biological processes of infected subjects. Furthermore, MYC activation is restricted to GSN, while these peptides in LNSN cannot overcome the P53 suppression and induction of cell death. Thus, unlike GSN, proliferation and angiogenesis are potentiated in LNSN, and the cell death mechanisms, including apoptosis and autophagy, are inhibited.

## Discussion

The main purpose of this study is to establish a system biology framework for understanding the principle features of *Mtb*, and its interactions with the host in the lung and local lymph node in the active TB, which in turn provides a new model-based vision for the development of cost-effective, safe and potent methods for the prevention and treatment of TB. The survival of *Mtb* as a parasite depends on host-microbe interactions, which can consequently lead to elimination, latency or acute TB [12]. Thus, in order to survive, *Mtb* has to escape from the host immune responses. It is suggested that *Mtb* is not a cytopathic microbe, and TB is an immunopathologic disease due to the host inappropriate immune response to *Mtb* colonization and the evasion of the microbe [13]. Therefore, the epigenetic mechanisms, such as the gene expression variations in host-*Mtb* interactions in a defined environment determine the outcome of this challenge [14]. However, the ecological niche of *Mtb* is a powerful cell of the immune system, i.e. the macrophages in the lung. According to the compartmental theory, the appropriate immune response must be developed in three different compartments, i.e. the lung, local lymph nodes, and blood circulation [15]. Therefore, in this study, the host-microbe interactions were assessed based on two high-throughput RNA sequencing (Seq) studies, which investigated the most important molecules implicated in the development and progression of TB. It is worth noting that the central molecules in such analyses can be targeted for the prognosis or treatment of these re-emerging life-threatening microbes.

With respect to the present analyses, although there are shared mechanisms in the lung and local lymph node responses, *Mtb* in particular, and the host in each compartment and each stage focus on their most powerful molecules that depend on the habitat, in which the challenge is in progress. The greatest challenge in these interactions in the granuloma compartment regarding *Mtb*, is fighting for survival by inducing the inappropriate immune response, while the host may attempt to induce the cell death in the infected macrophages or at least enclosed the site of infection with macrophage lining and fibrin formation [16]. *Mtb* in the lymph node attempts to hide and inhibit the host appropriate immune responses [17]. Overall, the main gene expression changes in the *Mtb-*infected cells in the granuloma is associated with the cell survival, proliferation and DNA repair, while the *Mtb* challenge in lymph node occurs in Ag presentation, inflammation, inhibition of apoptosis, autophagy, phagolysosome formation and Th1-Th17 differentiation [13, 18].

### Microenvironment manipulation

Caseous granulomas are typical compartment of TB, formed by epithelioid macrophages surrounding a cellular necrotic region, with a rim of T- and B-cell lymphocytes. However, once the granuloma is established, neutrophils do not seem to play an important role in the host responses, while neutrophils in caseous or TB recurrent stages are recruited to the site of *Mtb* dissemination [19]. Fig. 8 shows the signaling pathways that are manipulated by *Mtb* and form a micro-environment to respond.

As one of the most frequent cells in a granuloma, macrophages are the main effectors in killing *Mtb*, but they also act as the ecological niche for the *Mtb* replication [2]. According to the analyses in the present study, one of the survival targets for *Mtb* is the metabolic manipulation of the infected cells. Infected cells are proliferated in the caseum stage. Thus, the cell in such a situation needs to provide a new source of energy and more oxygen to survive in this new microenvironment [20]. Based on several studies, caseous necrosis is related to a hypoxic state and hypoxia signaling pathways, such as HIF1α pathway, which is induced [21, 22]. Moreover, some mechanisms including the angiogenesis and anaerobic metabolism are enforced by the elevation of MYC [23]. At the same time, the expression of EP300 and HDAC1 were down-regulated by negative feedback via acetylation and de-acetylation of the HIF1α downstream pathways in order to modulate this phenomenon [24, 25].

Furthermore, caseation is associated with an increase in lipid metabolism. Thus, the dominant nutrition changes to lipids and establishes the foamy macrophages in the formation of the central necrotic zone. According to the previous studies, the total serum triglycerides (TG) are higher than the normal and predominant storage lipids in the caseum and foam-cell-rich areas of tuberculous granulomas [26]. As P53 promotes fatty acid oxidation to support cell survival, a significant increase in the P53 expression in granuloma can be evidence of lipid uptake and metabolism in the signaling pathway of granuloma [27]. At the same time but unlike granuloma, the T cells, dendritic cells and infected macrophages in the lymph node, as the other typical compartment of the *Mtb* niche, are the main responding cells [28]. Fig. 9 demonstrates that the signaling pathways are manipulated by *Mtb* and form the microenvironment, which will be discussed in the following sections.

When macrophages or DCs engulf *Mtb*, they migrate from the lung as the compartment of infection to the regional lymph nodes in the T cell zone [29]. Forming the appropriate immune response in this compartment is very critical, and where specific *Mtb* CD4^+^ T cells recognize *Mtb* antigens bound to MHC molecules, the *Mtb*-specific T lymphocytes are activated and differentiate to the blast cell. These specific effector cells are then attracted to sites of *Mtb* replication via blood circulation as the third compartment [30].

The proliferation of T cells is induced to form the appropriate response against *Mtb*, so that the metabolism shifts into the anaerobic state [31]. In addition, angiogenesis provides more oxygen to survive in this new microenvironment of the lymph node. Similar to granuloma, the elevation of the MYC expression makes these two mechanisms of anaerobic metabolism and angiogenesis possible [23]. Lipid uptake and fatty acid metabolism are down-regulated in the lymph nodes to control the *Mtb* growth by changing the nutrition supply [32].

### Cell dynamics

It is suggested that apoptosis in the granuloma kills the intracellular bacteria and promotes Ag presentation, thereby stimulates the Th cells, whereas the necrosis of infected cells releases the bacteria and promotes the inflammation and tissue damage [33]. The tuberculosis necrotizing toxin (TNT) is a secreted nicotinamide adenine dinucleotide (NAD^+^) glycohydrolase that induces necrosis in the infected macrophages [34]. In addition, P53 induces necroptosis by transactivating the necrosis-related factor (NRF), cathepsin Q, and also directly binds with cyclophilin D (cypD) [35]. Simultaneously, P53 stimulates other non-canonical types of cell death, including caspase-independent apoptosis (CIA), ferroptosis, autophagic cell death, mitotic catastrophe, paraptosis, and pyroptosis [36].

Moreover, the overexpression of YWHAG, YWHAQ and NPM1are involved in apoptosis, leading to an increase of expression of BAG3 and VCP, which contributes to autophagy [37, 38]. Another mechanism that leads to apoptosis is the severe endoplasmic reticulum (ER) stress, which has induced inhibition of ubiquitination through the increase in COPS5, CAND1, OBSL and SIRT7, as inhibitors of the ubiquitin complex [39, 40]. Increasing HSPs as chaperone molecules confirms this ER stress. It is a smart mechanism that *Mtb* utilizes to prevent its proteins from degradation, antigen-presenting and hiding from the immune system [41]. Accumulation of lipid droplets in macrophages induced by *Mtb* mycolic acid promotes the associated microbe proliferation [42]. Furthermore, *Mtb* regulates the expression of host genes and induces cell proliferation. Elevation of MYC, CDK2, CUL3, YWHAG, MCM2, RPA2 and NF-κB in our study was involved in cell proliferation. In this regard, there is a dynamic condition between the cell components of granuloma, inducing the death or proliferation.

On the other hand, the following events occur simultaneously in the lymph node. It is assumed that apoptosis in the lymph node can induce T cell depletion or infected cell death that results in intracellular bacterial release and promotion of Ag presentation. Based on Fig. 9, apoptosis is inhibited by the overexpression of YWHAZ, YWHAB, CDK1, CDK2 and MYC genes and up-regulation of the NF-κB pathway, leading to the T cell survival in favor of the host. The overexpression of CDK1, CDK2, BRCA1, MYC and NF-κB is involved in cell proliferation in the signaling pathways. Therefore, there is cell dynamicity towards the proliferation and survival of the T cells as the main cells in this compartment, resulting in the up-regulation of the cell proliferation pathways and down-regulation of cell death pathways.

DNA repair is a crucial mechanism in the cell cycle and obviously cell proliferation. Therefore, the expression of genes involved in DNA repair should be elevated in parallel with proliferation pathways [43]. Although some molecules such as P53, CKD2, CDK1, UBC and MYC are playing in both activities, some others, e.g. CDC5L and FBXO6 play a selective role in the DNA repair in both the granuloma and lymph nodes (Fig. 8 and Fig. 9). It is worth to note that EP300, as the main player in epigenetic events, was down-regulated, which contributed to the DNA repair only in the granuloma [44]. According to Figs. 8 and 9, DNA repair is a crucial mechanism in the cell survival, and there are common gene expression changes in both the granuloma and lymph node compartments, while some others, including CDK1, TRAF6, ABL1 and BRCA1, are up-regulated in lymph nodes, selectively.

One of the determinative mechanisms of survival for the infected cells that occur in granuloma is the manipulation of epigenetic mechanisms, targeting HDAC1, EED, SIRT7 and EP300. Unlike acetylation that works as a pro-apoptotic mechanism, de-acetylation induces the survival [45]. The up-regulation of EED and SIRT7 as de-acetylation factors and down-regulation of EP300 as an acetyltransferase among these genes are beneficial for the *Mtb*, through reinforcement of the survival pathways. However, HDAC1 is down-regulated, which seems that the host is attempting to trigger apoptosis [24, 46].

Different studies showed that the *Mtb* infection up-regulates the *Sin3A* expression, and this gene is encoding a co-repressor that acts in concert with HDACs (histone deacetylases) to repress several genes, including MHC class II gene expressions [24]. Therefore, the down-regulation of HDAC1 is in favor of the host for promoting the Ag presentation [24].

However, gene expression manipulation by Mtb infection in the lymph nodes differs with granuloma, in which the survival mechanisms and proliferation are performed through the overexpression of EWSR1 and SIRT7. EWSR1, as a transcriptional repressor, disturbs the gene expression by mimicking or interfering with the normal function of CTD-POLII within the transcription initiation complex. This gene involves in the apoptosis inhibition and promotes proliferation and angiogenesis. In addition, SIRT7 acts as deacetylase and contributes to apoptosis suppression and proliferation [47, 48].

### Transcriptome manipulation

Another gene expression manipulation by *Mtb* in granuloma happens in transcriptome splicing, by down-regulating CDC5L, HSPA8 and SNW1 in Prp19 complex, and also up-regulation of HNRNPA1 and down-regulation of SRSF protein kinase 2 (SRPK2) in the common components. It is worth to note that CDC5L is the only splicing factor, also called E3 ligase. Thus, its down-regulation results in less ubiquitination, microtubule dysregulation and mitotic arrest, which may help to form giant cells [49]. SNW1 (SKIP) counteracts the P53 apoptosis pathway via selective regulation of p21 mRNA splicing. Thus, SKIP down-regulation inhibits both P53-mediated apoptosis in favor of the infected cell survival, and proliferation [50]. On the other hand, the HNRNPA1 overexpression, as an anti-apoptotic splicing factor of caspase-2, contributes to the inhibition of apoptosis, enhancing the expression of cyclin D and MYC by attachment to Internal Ribosomal Entry Site (IRES), and participating in splicing APP, leading to the induction of the NF-κB-mediated inflammation [51]. Moreover, HNRNPA1 stimulates telomere elongation and this is an important mechanism in favor of the proliferative cells. Furthermore, alternative splicing of IRF3 by HNRNPA1 leads to the reduction in IFN-β and pro-inflammatory cytokines [52]. It seems that *Mtb* suppresses innate immune mechanisms by manipulation of the HNRNPA1 expression. Another molecule, i.e. SRPK2, is also a highly specific kinase for the serine/arginine (SR) family of splicing factors, involving in apoptosis and differentiation of M1 macrophages. Hence, the low expression level of SRPK2 may inhibit apoptosis and increase the possibility of M2 macrophage differentiation, which left abandoned in caseum [53].

*Mtb* in the lymph nodes probably changes the splicing process by up-regulation of HNRNPA1 and down-regulation of CDC5L, in our analyses. This suggests that similar to granuloma, they involve in promoting proliferation and suppression of apoptosis in the lymph nodes [54].

### Proteome manipulation

Apoptosis and autophagy of macrophages play a pivotal role in the pathogenesis of TB [33]. In *Mtb* infection, autophagy can help macrophages to eliminate the bacterium. During autophagy, antigenic peptides are transported into autophagic compartments and degraded for peptide presentation; thereby, potentiating the protective cell-mediated immune responses [33].

In granuloma, the main *Mtb* strategies are attempting to prevent the infected cells from protein degradation and autophagy after that [2]. Ubiquitination is the main step of protein degradation; thus it is a strategic point for *Mtb* to prevent and resist against this strategy, targeting ubiquitin complex [55].

Indeed, increasing the expression of two gene sets has been observed in our analyses: (i) Components of ubiquitin complex such as CUL1, CUL3, FBXO6, HUWE1, Smurf1 and UBC; and (ii) Inhibitor of ubiquitin complexes like COPS5, CAND1, OBSL1 and SIRT7. The increase of inhibitors is probably another *Mt*b strategy, while the elevation of ubiquitin complex components is the host strategy in response to this suppression [55]. As a result, the misfolded proteins are increased, indicating a high expression of HSPs, which moderate the ER stress. Ubiquitinated proteins have three different destinations, i.e. entering to aggresome, autophagy or drive to proteasome or the immune-proteasome. The up-regulation of VCP and BAG3 shows an active role of aggresome that contributes to autophagy, which in this part could be the pressure of host responses. However, some studies report the up-regulation of Lamp2, Rab7, and Tfeb by PPAR-α in autophagy and lysosomal biogenesis as the host anti-mycobacterial responses [56]. In our RNA Seq analyses, only SMURF1 is up-regulated among the implicated factors in the autophagy, including LC3, LAMP1, and Smurf1. On the other hand, the proteasome is converted to immuno-proteasome by high amounts of IFN-γ and TNF-α induced by *Mtb*, but the Ag presentation is not efficient in the absence of ubiquitination [13, 57].

Same as granuloma, *Mtb* attempts to prevent the host from the degradation of its proteins in the lymph nodes, since a lymph node is the most significant compartment for the Ag presentation in order to establish an appropriate response, which is a strategic point for manipulation [13]. Ubiquitination inhibited by the overexpression of COPS5, CAND1 and SIRT7 contributes to the down-regulation of Ag presentation and autophagy [39]. Autophagy in the lymph nodes is related to Ag presentation by APCs, and hence both mechanisms have the same aim, i.e. inhibition of *Mtb* Ag presentation in favor of the microbe dissemination. Furthermore, suppression of autophagy increases the bacterial survival in the macrophages [13]. Ubiquitination is also manipulated by increasing the expression of two gene sets: (i) Components of ubiquitin complex such as CUL1, FBXO6 and UBC, and (ii) Inhibitor of ubiquitin complexes such as COPS5, CAND1 and SIRT7. The increase of inhibitors is probably the *Mtb* strategy, while the elevation of ubiquitin complex components is the host strategy in response to this suppression [2]. As a result, the misfolded proteins are increased and its reflection is the expression of HSPs, which moderate the ER stress. To conclude, three consequences of protein ubiquitination including the aggresome and autophagy, proteasome and protein degradation, and immune-proteasome and Ag presentation are down-regulated due to the suppression of ubiquitin complex [2].

### Host immune response and *Mtb*

The complex host immune responses, which are necessary for the *Mtb* elimination might be a Th1-dominated pattern, producing IFN-γ and IL-2, which can recruit and activate macrophages and CD8^+^ CTLs along with mild regulatory Th2 and regulatory T (Treg) cell responses to prevent the pathological damage [58]. Such a complex mechanism might be the immunological basis for the TB prophylaxis and treatment. In this regard, type IV hypersensitivity overexpresses IFN-γ as a pro-inflammatory cytokine. On the other hand, our analyses show that an increase in expression of UBE2I or NEMO as a SUMOylation factor plays an important role in promoting inflammation through the NF-κB pathway [59, 60]. Hiding from the immune system is always the first mechanism that *Mtb* chooses. In this study, the decrease in exportin 1 (XPO1) expression in both granuloma and lymph nodes is observed, which leads to the inhibition of nuclear export of HLA-A mRNA and IFN-I mRNA, and consequently, the Ag presentation to CTLs and immune response potentiation are suppressed by interferon type 1. Furthermore, the SMURF1overexpression suppresses the TGF-β pathway, leading to inhibition of the differentiation of T cells to Treg [55]. On the other hand, increasing the P53 level but not toward p21 activation contributes to the expression of Foxp3 and STAT5, which promotes Treg cell differentiation. Therefore, as the main producer of TGF-β, Treg can act as a double-edged sword in the TB manifestation, as it modulates the *Mtb* Th1 immune response to prevent type IV hypersensitivity, and it also prevents tissue damage [61]. Thus, modulation of the TGF-β pathway has a narrow range in favor of the host to help the formation of fibrosis around the infected site, but at the same time, it can suppress CMI in favor of the *Mtb* dissemination [61]. In contrast, some studies demonstrated that TGF-β may be involved in the tissue damage and fibrosis during TB, as it promotes the production and deposition of macrophage collagenases and collagen matrix [62]. Several studies demonstrated that *Mtb* induced HDAC1, leading to the inhibition of IL-12 and Th1 differentiation to form CMI, but HDAC1 in these analyses was decreased in a granuloma, which may act as a suppressing event in favor of the *Mtb* dissemination [24].

Some important events only occur in lymph nodes and not in granulomas, such as down-regulation of CALM and EGFR, both of which contribute to inhibition of phagolysosome formation, nitrite oxide (NO) production, and Th1/Th17 differentiation. The CALM and EGFR down-regulation can also inhibit Ag presentation by preventing the fusion of endocytosis vacuole to the endosome [48]. Certain studies have suggested that the inhibition of the iNOS expression and NO production can be considered as a mechanism for the propagation of various infectious agents, such as *Mtb*. Therefore, suppression of NO production in this study, by down-regulation of CALM and EGFR, can induce tissue damage resulting in elevated levels of MMP9.

The suppression of the first step of establishment of the effective adaptive immune response i.e. Th immune responses, mainly the upstream of epigenetic events toward Th1 differentiation, should be a promising target in the prophylaxis of TB. Furthermore, inhibition of both of the key mechanisms of Ag presentation and Th1 differentiation seems to be the main manipulated point in the lymph nodes. As a result, the inhibition of Th17 in parallel with Th1 guarantees that IL-17 cannot provide IL-12 Th1 differentiation.

A proper modulated immune response requires an appropriate regulation, which mainly forms by Treg and Th2 responses to inhibit the hypersensitivity reactions, and prevents immuno-pathological injuries [58]. In this study, a decrease in the SMAD expression, involving in the TGF-β pathway, leads to suppression of T cell differentiation to Treg cells. Unlike granuloma, the up-regulated P53 is inhibited by the upstream pathways, which cannot contribute to promoting the Treg cell differentiation by expression of Foxp3 and STAT5. Following the infection, mycobacterial products such as Lipoarabinomannan (LAM) induce the production of TGF-β by monocytes and DCs at the site of disease. It seems that *Mtb* attempts to induce more TGF-β production, while, the host performs the down-regulation to moderate this cytokine.

## Conclusion

Although in this method of system microbiology analyses, there are shared activities in different compartments of bacteria-host conflicts, there are selective events that also occur in those compartments. For example, the most considerable molecular expression and pathways affected by *Mtb* in granuloma are associated with monocyte/macrophage in the lung, while most of the changes in the suppression or activation in local lymph nodes are related to the T cell sub-populations. Based on the compartmental theory, the appropriate responses are important in bacterial elimination and disease manifestation. For example, in the current study regarding the TB patients, the RTK (EGFR) pathway, which is important in T cell activation and Th1 and Th17 differentiation, is down-regulated. Interestingly with this suppression, the Ca^2+^ dependent calmodulin molecule and consequently, NFAT formation and Erk/MAPK pathway are affected, and the T cell activation and differentiation are inhibited. More importantly, MYC, NPM1 and MDM2 in granuloma activate P53 in different ways and support the monocyte/macrophage survival in favor of *Mtb* and its dissemination, while PI3K pathway inhibits P53 in lymph nodes in the absence and suppression of NPM1. However, despite the up-regulation of proliferation from different pathways, *Mtb* attempts to inhibit DNA damaging, which kills its habitat by overexpressing the BRCA1, CDK1 and BCR/ABL in the lymph nodes, as well as FBXO6, CDK2 and CDC5A in the granuloma and lymph node compartments. The survival pathways such as NF-κB and PI3K/Akt also increase the inflammation reactions; for example, the overexpression of TRAF6 in the lymph nodes, and IKKs and UBE2I in granuloma contributes to the NF-κB inflammatory pathway. In both compartments, the antigen releases induce inflammatory reactions, but the Ag presentation is suppressed through the down-regulation of XPO1 and suppression of ubiquitination. However, overexpression of VCP, BAG3, Smurf1 and YWHAZ in the granuloma induces autophagy to support the Ag presentation and cell death, although it is suppressed by the mTOR signaling in the lymph nodes. In addition, the cell proteasome with the degradation of *Mtb* proteins supports more suppression of Ag presentation in the lymph nodes. It seems that these mechanisms in granuloma in the TB patients induce more inflammatory reactions in the form of type IV hypersensitivity, and consequently result in lung damages and inappropriate responses in the lymph nodes.

Although the granuloma is located in the lung, hypoxia and angiogenesis are more significant in this compartment than the lymph node, which is suppressed through the inhibition of HIF1-α via the down-regulation of EP300 and HDAC1. HIF1-α is an activator of angiogenesis, while in a granuloma, through the down-regulation of EP300 and HDAC1, HIF1-α is inhibited and functions as a suppression signal in this compartment. This comprehensive study may confirm the compartmental theory and also show the main targets for immune therapy. More studies are required with more validated methods to specify the exact mechanism of combating TB as a re-emergent life-threatening drug-resistant disease.

## Acknowledgments

The authors have a great thanks to our colleagues in Inflammation and Inflammatory Diseases laboratory, in particular, Ms. Narges Valizadeh, Ms. Saeedeh Mehraban and Ms Sanaz Ahmadi for their helps and supports.

